# Optokinetic response in *D. Melanogaster* reveals the nature of common repellent odorants

**DOI:** 10.1101/2024.05.17.594662

**Authors:** Giulio Maria Menti, Matteo Bruzzone, Mauro Agostino Zordan, Patrizia Visentin, Andrea Drago, Marco Dal Maschio, Aram Megighian

**Affiliations:** Padova Neuroscience Center, Università degli Studi di Padova, Padova, Veneto, Italy; Department of Biomedical Sciences, Università degli Studi di Padova, Padova, Veneto, Italy; Department of Neuroscience, Università degli Studi di Padova, Padova, Veneto, Italy; Department of Biology, Università degli Studi di Padova, Padova, Veneto, Italy; Entostudio S.r.l., Ponte San Nicolò (PD), Veneto, Italy

**Keywords:** Drosophila melanogaster, Repellents, Optokinetic response, Navigation

## Abstract

Animals’ ability to orient and navigate relies on selecting an appropriate motor response based on the perception and integration of the environmental information. This is the case, for instance, of the optokinetic response (OKR) in *Drosophila melanogaster*, where optic flow visual stimulation modulates the walking or flying patterns. Despite a large body of literature on the OKR, there’s still a limited understanding of the impact on OKR of concomitant, potentially conflicting, inputs. To evaluate the impact of this multimodal integration, we combined in *D. melanogaster*, while flying in a tethered condition, the optic flow stimulation leading to OKR with the simultaneous presentation of olfactory cues, based on repellent or masking compounds typically used against noxious insect species.

First, this approach allowed us to directly quantify the effect of several substances and their concentration, on the dynamics of the flies’ OKR in response to moving gratings by evaluating the number of saccades and the velocity of its slow phase. Subsequently, this analysis was capable of easily revealing the actual effect, i.e. masking vs repellent, of the compound tested. In conclusion, we show that *D. melanogaster*, a cost-affordable species, represents a viable option for studying the effects of several compounds on the navigational abilities of insects.

## Introduction

The navigational abilities of insects are remarkably close, if not equal, to the skills exhibited by animals commonly addressed, incorrectly, as “more evolved” or, better, as more complex. Meanwhile, the debate is still ongoing, asking whether these notable similarities may be identified as analogies due to the convergent evolution of independent nervous systems (Farries, 2013; Moroz, 2009; Northcutt, 2012), rather than reflecting homologies originating from the last common ancestors between Invertebrates and Vertebrates (Strausfeld and Hirth, 2013). The study of insects’ navigation flourished since the end of the 19^th^ century (Hect and Wald, 1934), resulting nowadays in extremely refined studies from the anatomical, physiological, engineering, and physics point of view. Nevertheless, at present, unravelling the mechanisms of insects’ navigation is becoming of even more importance for tackling the challenges placed by some species to agriculture and human health.

In Europe, the increasing presence of alien insect species, exacerbated by the occurring variations in climate, is posing a serious threat to cultivation, as is the case for the dipteran *D. suzukii* from Southeast Asia (Tait et al., 2021), and to public health, which is endangered by several vector-transmitted diseases spread by insects like mosquitoes and ticks (Semenza and Suk, 2018). Moreover, in 2012, the approval of the EU Biocidal Product Regulation thinned the list of compounds available for companies and consumers to distribute and purchase (Regulation (EU) No 528/2012). Regarding the public health aspect, and particularly the threat posed by mosquitoes (Giunti et al., 2023), a deeper comprehension of the effects of the repellent compounds still publicly available for production and commercialization could increase the efficacy of repellent products and enhance the global protection against vector insects in the population. However, identifying a suitable approach to study these aspects is still challenging: the rearing of mosquitoes and other pest species requires special attention and carefulness (see the latest 2021-22 FAO/AIEA guidelines), which are not always easy to satisfy in public laboratories with limited spaces and resources. In turn, *D. melanogaster*, a well-known model organism, does not require extensive care, especially when dealing with wild-type strains, and it also shares with mosquitoes the same brain organization and the sensitivity to some known mosquitoes’ repellents. Reports in at least one other species in this *genus* (*D. suzukii*) (Afify et al., 2019; Syed et al., 2012) have been confirming this view.

Therefore, we considered whether *D. melanogaster* could be a suitable organism for investigating both the raw repellent effect of the substances of choice, and their possible influence on the navigational abilities of insects This strategy would open the possibility to utilize the extremely sophisticated behavioural, neurophysiological, and genetic techniques available in this model organism, using it as a probe organism in screening for the effects of promising repellent compounds. However, the ecological differences between different insect species should always be accounted for, as one compound could be in theory repellent to one species but not to another or have opposite effects for larval and adult stages.

We decided to tackle these questions by setting up a behavioural paradigm with adult *Drosophila* in fixed tethering conditions, where the tethered fly is not able to rotate around the vertical axis. We investigated the OKR of these flies presented with a moving grating visual stimulus (Cellini et al., 2021; Götz, 1964; Hect and Wald, 1934) during the concurrent administration of different compounds chosen among the ones already being tested on mosquitoes, namely: eugenol, lemongrass, picaridin, and IR3535^®^ (Afify et al., 2019).

Our research extends the body of literature investigating the OKR in flies or the effects of odorants on their flight conducted in the past years (Chow and Frye, 2008 and 2009; Duistermars et al., 2009), as well as studying the multisensory integration in flies’ navigation (Currier and Nagel, 2020), for instance, on vision and olfaction in presence of attractive odour cues (Duistermars and Frye, 2010). As readout, being the fly unable to turn, we chose to track the dynamics of the orientation of the head, focusing on the movements elicited by the optokinetic stimulation. In flies, this stimulation elicits nystagmus-like sawtooth head oscillations, which closely resemble the typical optokinetic nystagmus (OKN), observed for eye movement in other animals, humans included. These nystagmus-like head movements, which herein we will refer to as “head optokinetic nystagmus” (HOKN) are characterized, as in the classical OKN, by two alternate phases, a slow and a fast phase, taking place when the visual system is properly stimulated. The slow phase, consensual to the movement of the visual panorama, encapsulates the reflexive response to the motion of the visual field (or visual stimulus) and is generated by the optokinetic neuronal circuits. This slow phase is immediately followed by a second fast phase, identifiable as a rapid ‘reset’ movement of the head in the direction opposite to the moving panorama (Cellini et al., 2021). Said fast phase happens as the flies, while following a moving panorama, come closer to the torque limits of their heads and need to reset their gaze to keep pursuing the shifting visual stimulus (Cellini et al., 2021). This resembles the eye dynamics found in humans and mammals, where the fast phase is generated by resetting circuits located in the pontine reticular formation for the horizontal OKN. (Land, 2019).

In our study, we looked for the differences in the dynamics of the HOKNs during the OKR of flies exposed to a plume flowing in the opposite direction of the visual stimulus and carrying a repellent compound. Our working hypothesis was that if the repulsive cue evoked by the odorant had been strong enough, this would have interfered with the dynamics of the elicited HOKNS. We found that aversive compounds can affect the OKR of flies, decreasing the number of HOKNs and increasing the delay in between each event, while leaving the intrinsic dynamics of the process (e.g. the angular velocity during the slow phase) unaltered.

Moreover, we think our findings demonstrate that this kind of approach is feasible, being relatively simple to set up, and affordable. Moreover, it could be extended to other *Drosophila* species or, with adequate adjustments, other insects, being a *trait d’union* between studying their navigational abilities and the necessity of testing suitable products for both agriculture and health care.

## Results

### Measuring HOKN in flies

To investigate the effect of multimodal sensory integration on the fly optokinetic response, we started by identifying the relevant kinetic parameters of the head movements associated with the presentation of a visual stimulation based on horizontally moving grating (Fig. 1A). We considered that the head movement represents a good proxy for the OKR response, presenting an OKN-like pattern with a slow turning phase in the same direction of the grating motion, followed by a fast reset phase in the opposite direction (Fig. 1C). We quantified these responses, using an automated behaviour analysis pipeline to extract the total number of head saccades during the stimulation period, the characteristic inter-saccadic interval, and the corresponding angular velocity for the slow phase. For the considered grating movement speed (60 deg.sec^-1^), the acquisitions presented a generally consistent pattern across the different flies tested (good vs discarded), confirming that this kind of head movements or head saccades can be considered a reliable readout of the OKR response.

### Design of the multimodal sensory input protocol

Along with the visual stimulation, we designed a protocol to assess the impact of a multimodal sensory integration on the HOKN features, delivering an odorant flow directed opposite to the grating movement and with either neutral odour (plain mineral oil solution) or odorant compounds (Fig. 1A). The compounds were chosen considering two odorant types, a group of two natural repellents, namely eugenol and lemongrass oil, along with a group of synthetic compounds, picaridin, and IR3535^®^ (Fig. 1B), classified according to reported test on mosquitoes; this classification considers the observed mechanisms of action for the two groups, as the natural substances have an active repelling effect, while the synthetic ones act as masking agents which reduce the volatility of molecules relevant to the insects’ antennae (Afify et al., 2019). All the compounds were diluted in mineral oil and tested with two concentrations (0.5% and 1% in volume). Groups of flies, all naive to the odorants, were tested for a single odorant at a single concentration. We designed a multimodal integration protocol with an initial time segment (40 seconds) where flies were exposed to visual stimulation with neutral airflow. This consisted of a fixation phase (Buridan, 10 secs) followed by drifting gratings (Mineral Oil Phase, MOP, 30 secs). In the following segment visual stimulation was paired with the odorant stimulation (Odorant Phase, OP, 30 secs) during the grating motion phase. This same trial was repeated six times separated by a 20 seconds (Pause) dark period (Fig. 2).

**Figure 1.**
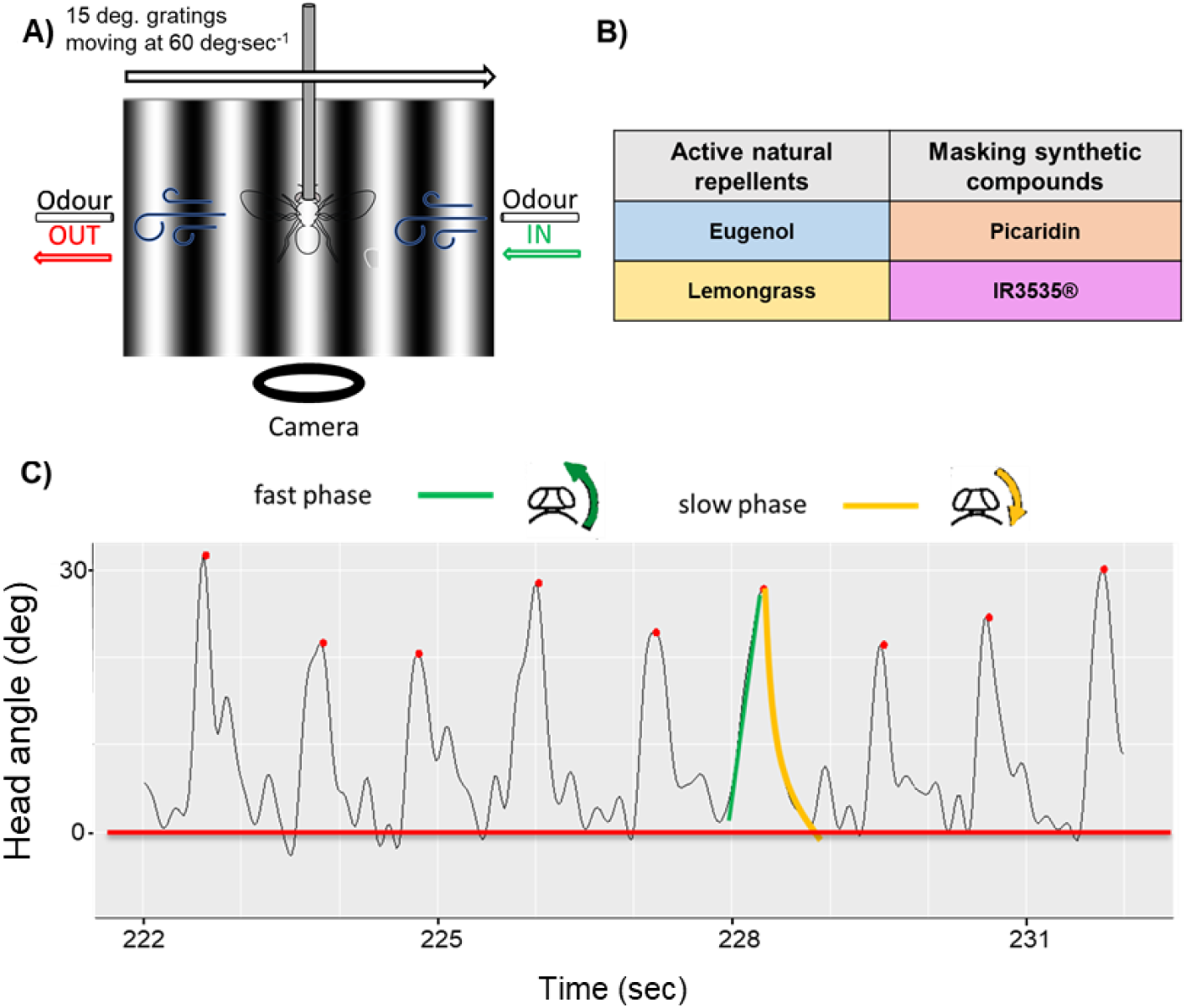
Experimental paradigm. A) Setup scheme: the whole apparatus is placed in a dark chamber and the illumination is achieved through infra-red LEDs. The fly gets its thorax glued to a pin and is placed on a fixed support (unable to rotate around the z-axis). A screen for projecting the visual stimuli (gratings pattern drifting to the right of the animal at 60 deg.sec^-1^) is placed in front of the fly, which is suspended inside a continuous air flow going in the opposite direction with respect to the visual stimulus. The camera for recording is placed below the animal. **B)** Tested compounds: we chose 4 substances classified by their nature and known mechanism of action (Afify et al., 2021). Two compounds are natural repellents with an active repulsing effect on insects (eugenol and lemongrass). The other two are synthetized substances (picaridin and IR3535®) with a masking action mechanism, binding and making less volatile other molecules which could be relevant cues to the insects. **C)** Saccade scheme and identification: we based our analysis on the flies’ reset head-saccades. As the flies experience and follow the optic flow (OF) generated by the frontal drifting visual stimulus, their head reach its angular limit and a reset saccade occurs, allowing the fly to keep pursuing the OF. Saccades are characterized by a fast phase (green), in the opposite direction of the moving visual stimulus, followed by the slow phase (gold), concordant with the OF. We identified the saccades by tagging the ‘peaks’ in the raw tracks of the head position with the ‘Findpeaks’ function in MATLAB and extracted the velocities of the slow phase ((peak location + 18 frames) / time) and the fast phase ((peak location – 5 frames) / time, not analysed).

**Figure 2.**
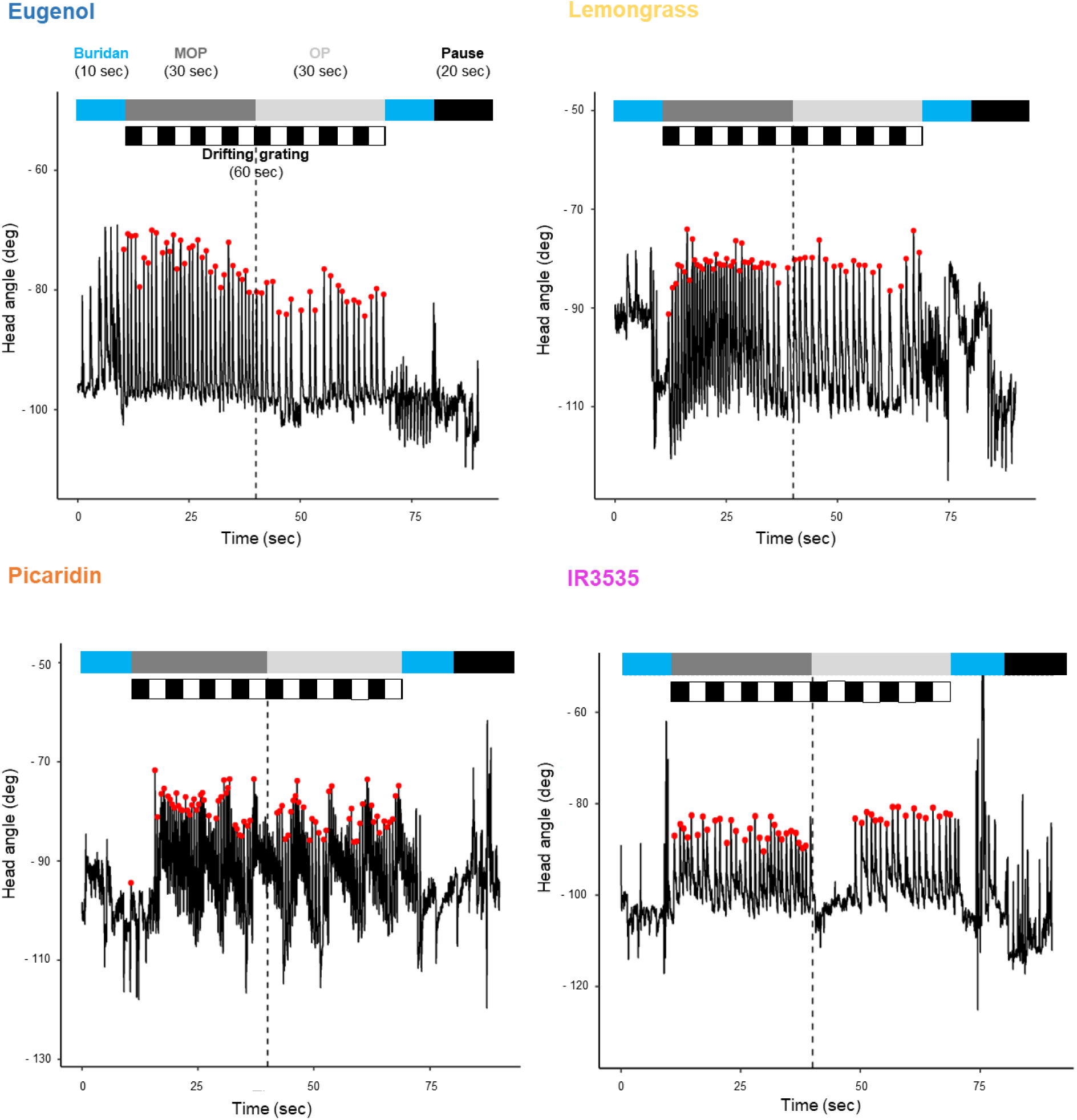
Raw recordings. Zoom of the 4th trial for one track from each group: each track shows a visible change in the number and/or inter-saccadic intervals in between events, as reported further down the result section, during the Mineral Oil Phase (MOP), when no odour is present, or the Odorant Phase (OP), when repellents are delivered to the fly. The shown paradigm was repeated six times, for a total of 600 seconds in each experiment (one fly). Y axis values are shown as negative as per the original FlyAlyzer output: the -90 angle corresponds to a fly’s straight head. The corresponding full tracks are plotted in Supplementary Figure 1.

### Repellents do not alter the saccade intrinsic kinetics

Even though the olfactory processing channel is expected to not directly impact the neuronal components in charge of the saccade motor execution, we wondered whether the presence of a repellent gradient opposed to the optic flow direction could impact the tracking process of the fly. Therefore, to check this first hypothesis, we started our analysis focusing on the velocity of the slow phase, when the head of the insect, following the optic flow direction, moves toward the odour source. Comparing this parameter between MOP and OP, in general, we did not find any significant difference (Fig. 3 and Table 1). The only exception from this generalized trend is represented by the condition with lemongrass at 1%, showing a significant reduction in the slow-phase velocity.

**Table 1.**
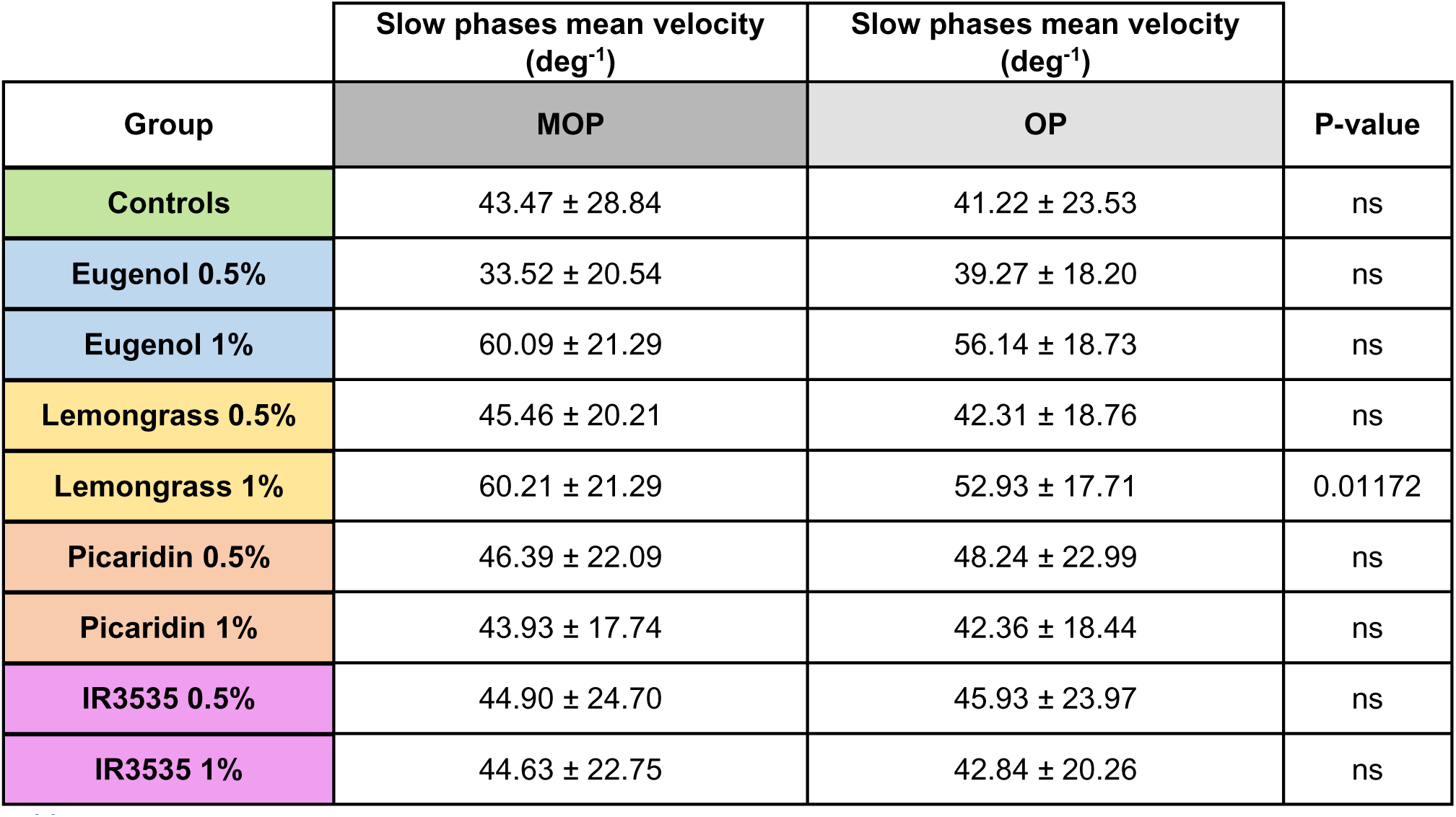

**Figure 3.**
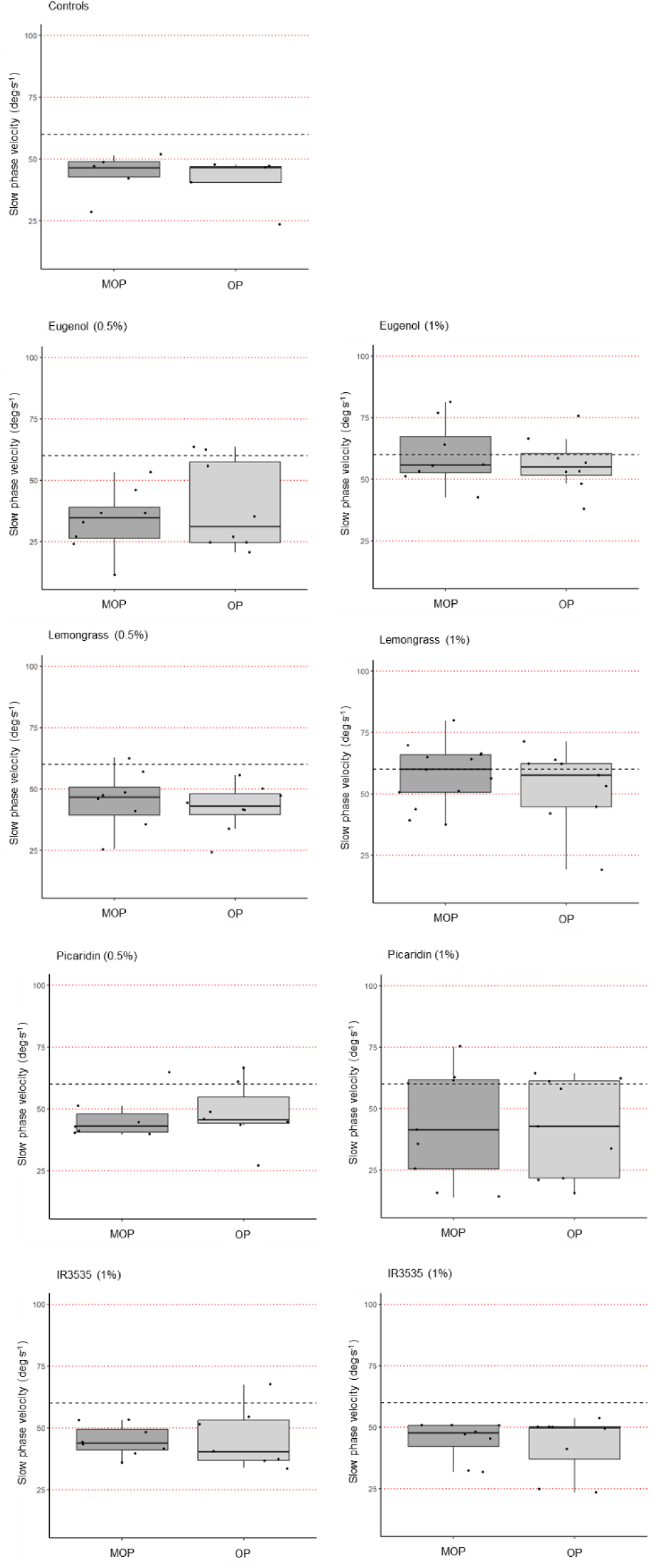
Slow phase velocities. Standard boxplots showing the median slow phase velocities mediated by subjects between MOP and OP in each group (limits of the box mark the 25th and the 75^th^ percentile). The dashed horizontal line is the reference for the grating pattern velocity. The only meaningful difference is observed (see Table 1) within the L1 group. Statistical analysis was performed checking for normality and heteroscedasticity and then applying the Mann - Whitney U test.

### Odorant presentation induces a decrease in the HOKN number

To further evaluate the possible effect of the odorants, we started our analysis focusing on the number of HOKNs and their relative change in numerosity from MOP to OP, with respect to the corresponding values measured in the control group, exposed only to mineral oil across the whole protocol (fake OP). We observed a generalized trend for the natural repellents, with a lower number of OP saccades in all groups and concentrations when compared to MOP: [**Controls** (**C**)] mean MOP 169.40 ± 89.80 vs mean OP 175.60 ± 50.71 saccades (103.65%); [**Eugenol 0.5%** (**E05**)] 124.62 ± 62.77 vs 91.50 ± 29.56 (73.42%), p = 3.326e^-7^; [**Eugenol 1%** (**E1**)] 201.46 ± 83.02 vs 160.06 ± 87.67 (79.45%), p = 1.149e^-6^; [**Lemongrass 0.5%** (**L05**)] 136.87 ± 33.41 vs 120.87 ± 40.53 (88.31%), p = 0.00862; [**Lemongrass 1%** (**L1**)] 188.78 ± 43.84 vs 143.57 ± 51.26 (76.04%), p = 3.232e^-8^. The effect was significant at both the concentrations tested with no statistically significant evidence for a dose-dependent effect, except between **L05** and **L1** (p = 0.0056). Interestingly, for the synthetic repellent, a dichotomic pattern emerged, with Picaridin showing a decrease in the number of saccades: [**Picaridin 0.5%** (**P05**)] mean MOP 160.57 ± 75.91 vs mean OP 117.28 ± 32.42 saccades (73.04%), p = 1.027e^-7^; [**Picaridin 1%** (**P1**)] 152.60 ± 65.49 vs 119.50 ± 60.57 (78.30%) p = 3.918e^-6^, similarly to natural repellents, and IR3535^®^ resulting the only odorant not affecting the number of OP saccades: [**IR3535 0.5%** (**I05**)] 177.87 ± 108.99 vs 175.75 ± 73.34 saccades (98.80%); [**IR3535 1%** (**I1**)] 107.87 ± 72.95 vs 105.50 ± 70.77 (97.79%).

**Figure 4.**
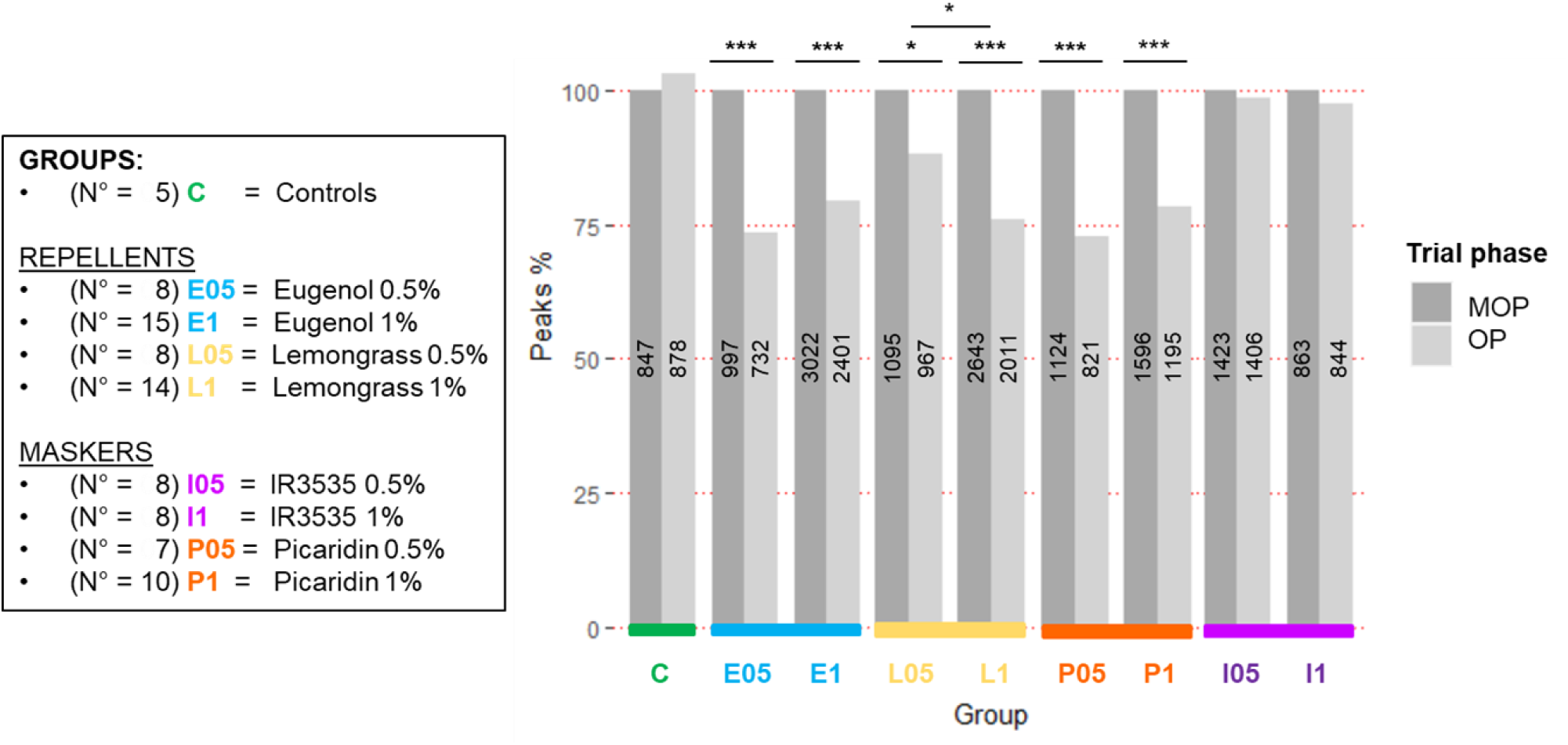
Saccades (peaks) number and proportion. The bar plot shows the relative proportion of MOP and OP HOKNs normalised on the MOP; values within each bar refer to the total number of HOKN identified during each phase in each group. P-values result from multiple 2-sample tests for equality of proportion with continuity correction, to verify if the observed differences in proportions between MOP and OP differed significantly from the **C** group and between the two concentrations within each group. *** : p < 0.001, ** : p < 0.01, * : p < 0.1.

### Inter-saccadic interval increases in the presence of the repellent

Focusing on the origin of the reduction in the number of saccades, we considered two scenarios. The first one, where the input from the olfactory channel, stimulated by the repellent, could with time overtake the HOKN mechanism, leading to a progressive decrease in the number of the saccades. The second one is where the visual and the olfactory channels co-exist, and the repellent interference does not suppress the normal HOKN but rather modulates the frequency of the saccades. We then looked at the Inter-Saccadic Interval (ISI), the time interval separating two consecutive saccades, comparing this parameter during MOP and OP (Fig. 5).

As expected, the control group, exposed only to mineral oil, did not show any appreciable difference in the ISI), showing a substantial overlap of the corresponding distribution in MOP and OP (2W-ANOVA [F = 35.338, num. d.f. = 17.0, denom. d.f. = 7451.4, p-value < 2.2e^-16^] + post-hoc Games-Howell test). Things were different for the repellents, as visible in Figure 3, where MOP and OP distributions of the ISI in both concentrations are shown for each compound against the **C** control distribution (black line).

As for the natural repellents, eugenol and lemongrass, the distributions are characterized by a significant increase in the ISI for most of the concentrations (Fig. 5A and 5B): **E05**, mean 1.18 ± 1.53 vs 1.61 ± 1.63 sec. (+36%, p = 5.9712e^-6^); **E1**, 0.76 ± 0.8 vs 1.14 ± 0.92 sec. (+50%, p = 1.2536e^-6^); **L05**, 1.13 ± 1.28 vs 1.27 ± 1.42 sec. (+12.5% [ns]); **L1**, 0.77 ± 0.84 vs 1.00 ± 1.10 sec. (+30%, p = 6.7175e^-9^). Conversely, in the case of the synthetic compounds (Fig. 5C and 5D), the scenario reflects the data related to the number of peaks, with no significant differences in the ISI distribution for IR3535^®^ at both tested concentrations: **I05**, 0.89 ± 1.3 vs 0.87 ± 1.00 sec. (-2.25% [ns]) and **I1**, 1.20 ± 1.67s vs 1.22 ± 1.58 sec. (+1.6% [ns]). Picaridin, on the other hand, shows features more like those observed in natural repellents also in the ISI rather than more similar to the other synthetic masking agent, with a clear increment in the mean ISI at both concentrations (**P05**, mean 0.87 ± 1.01 vs 1.30 ± 1.92 sec. (+49%, p = 1.2860e^-6^); **P1**, 0.94 ± 1.18 s vs 1.18 ± 1.43 sec. (+25%, p = 0.0004)).

**Figure 5.**
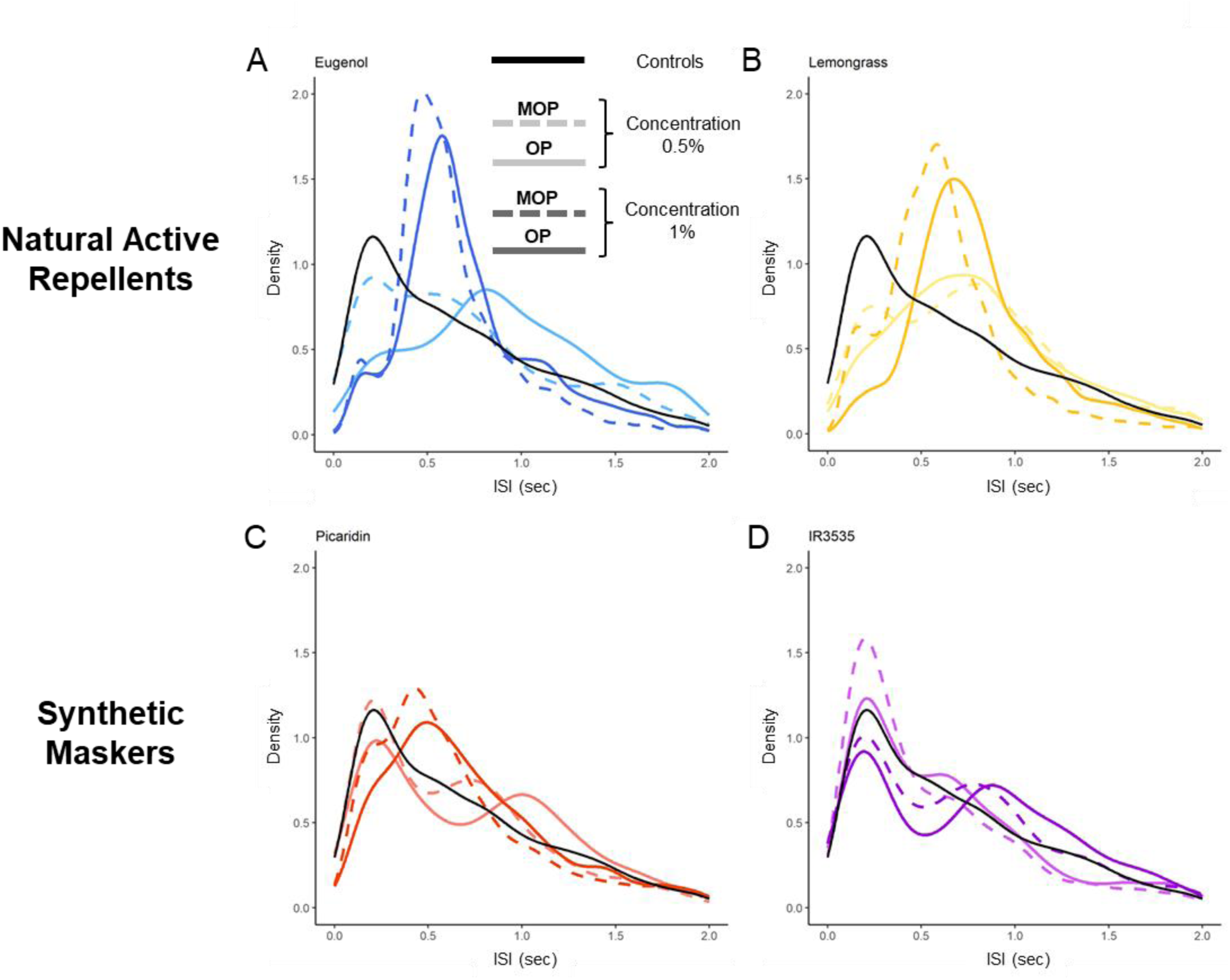
ISI distribution. The plots show the distribution of the ISI in the four groups, following the same colour scheme from the above figures. The lesser concentrations are represented by the lighter colours, and the MOP and OP are shown as dashed or continuous lines, respectively. The continuous reference black lines represent the whole Control group’s ISI, unsplit into MOP and OP since there was no statistically significant difference between the two. Statistical analysis was achieved through a 2W-ANOVA followed by a post-hoc Games-Howell test (the relevant p-values are reported in the main text ‘Inter-saccadic interval increases in the presence of the repellent’ section).

### Inter-Peaks Interval duration seems to slightly reduce as trials proceed

Additionally, we noticed a trend while looking at ISI value variations across trials in the groups (see Supplementary Fig. 2). As shown in Fig. 6, there is a subtle trend towards the ISI reduction as the trials proceed (1^st^ to 6^th^) and it was detectable in both the MOP and the OP. The mean values and standard deviations are shown below (Table 2). We first evaluated this trend applying a simple linear model (LM), and we found a slight but significant effect on the ISI with the increasing trial number in all groups (**C**: p **=** 2.00e^-9^; **E05**: p = 2.84e^-11^; **E1**: p = 5.50e^-11^; **L05**: p = 0.035; **L1**: p = 9.21e^-7^; **P05**: p = 1.03e^-12^; **I05**: p = 0.00248; **I1**: p < 2e^-16^) except for **P1**. Then we tested the mean values of ISI in the 1^st^ trials against the 6^th^ and found a significant reduction of the ISI, both during MOP and OP (see Table 2), in **C** (MOP: 2.21 ± 1.91 vs 1.06 ± 0.71 sec.) and **I1**(MOP: 3.4 ± 2.56 vs 1.64 ± 1.16 sec.; OP: 3.4 ± 3.5 vs 0.96 ± 0.48 sec.), and also during the OP in the 1% concentrations of both **E1** (OP: 0.91 ± 0.29 sec.), and **L1** (OP: 1.7 ± 1.26 vs 0.96 ± 0.22 sec.). Moreover, returning back to the underived data (the number of HOKN per second in each group), we normalized them for the number of flies in each group, checked if there was a meaningful difference in this variable through groups in our original dataset (Kruskal-Wallis test, chi-squared = 476.83, d.f. = 8, p < 0.001 and Dunnet test with Bonferroni correction for pairwise comparisons, see Table 1 in Supplementary materials), and we also checked through a permutation test of 10^4^ iterations that the observed differences could not be replicated by randomly re-assigning the observations within the groups (p < 0.001). Then we looked for the best fitting from different generalized linear models on the collective data (not separated for MOP or OP, since again there was no significant difference in their distributions) from **C** and **I05**, which did not show significant differences. We found the best fit to be a second-order polynomial regression, with the following R-squared values: **C**, 0.10596417; **E05,** 0.10085333; **E1,** 0.02766739; **L05** 0.02097825; **L1**, 0.15916139; **P05**, 0.14127738; **P1**, 0.04623636; **I05**, 0.05301220; **I1**, 0.27136975.

We also calculated an “Effect Index (E.I.)” for the ISI differences observed over time matching the OP and MOP from their corresponding groups:

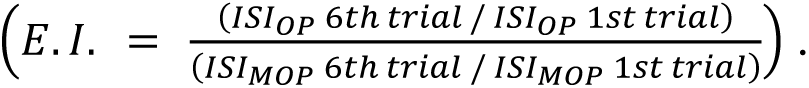

This index describes ideally the possible strength each compound could have in perturbing this slight decline in ISI (Table 3): scores close to 1 (**C**, **L05**, or **I05**) indicate that the diminution in the ISI was even and not much affected by the presentation of compounds, while scores lower than 1 (**E1**, **L1**, etc.) indicate that the major reduction happened during the MOP (repellents effectively impaired said reduction during the OP), and conversely, one score way greater than 1 (**P1**) shows that the change occurred mostly during the OP.

**Figure 6.**
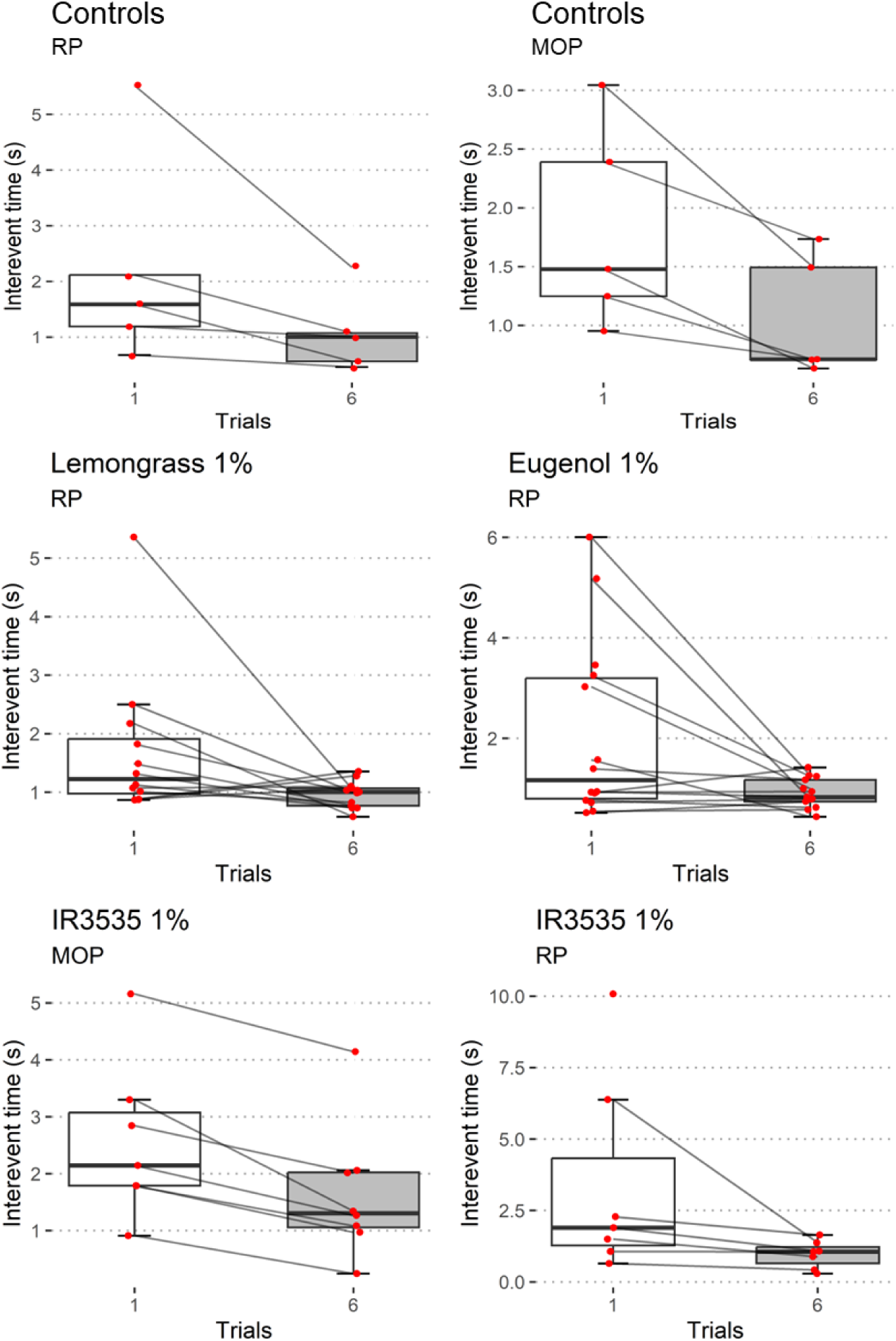
ISI mean values between the 1st and 6th trials. Standard boxplots representing the median ISI values divided by trial and mediated per subject (only statistically significant comparisons from Mann - Whitney U tests are shown).

**Table 2.**
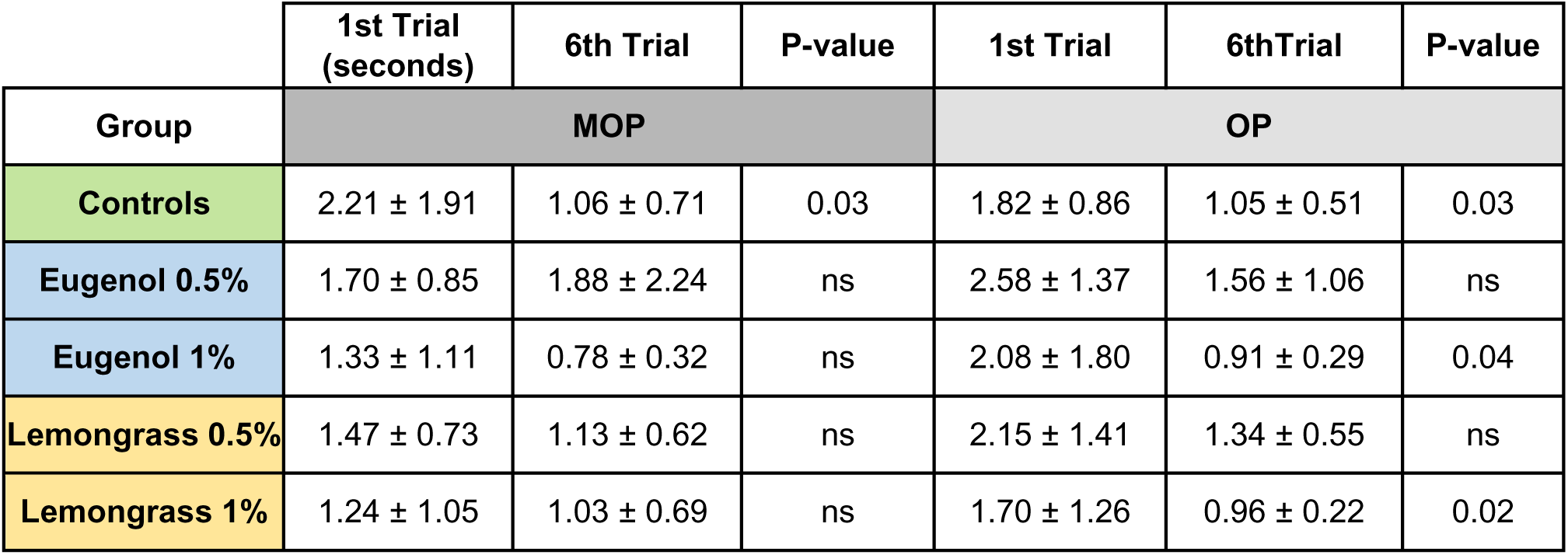
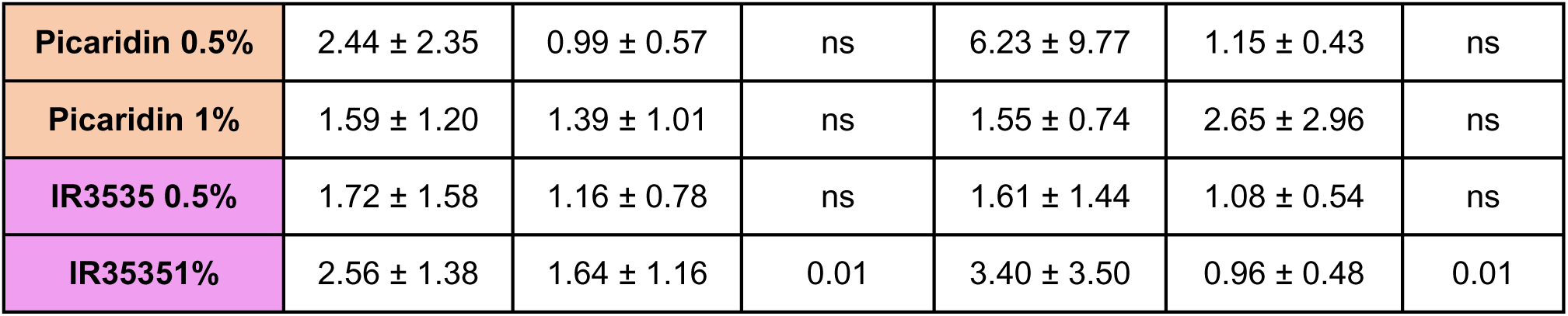

|  | 1st Trial<br>(seconds) | 6th Trial | P-value | 1st Trial | 6thTrial | P-value |
| --- | --- | --- | --- | --- | --- | --- |
| Group | MOP |  |  | OP |  |  |
| Controls | 2.21 ± 1.91 | 1.06 ± 0.71 | 0.03 | 1.82 ± 0.86 | 1.05 ± 0.51 | 0.03 |
| Eugenol 0.5% | 1.70 ± 0.85 | 1.88 ± 2.24 | ns | 2.58 ± 1.37 | 1.56 ± 1.06 | ns |
| Eugenol 1% | 1.33 ± 1.11 | 0.78 ± 0.32 | ns | 2.08 ± 1.80 | 0.91 ± 0.29 | 0.04 |
| Lemongrass 0.5% | 1.47 ± 0.73 | 1.13 ± 0.62 | ns | 2.15 ± 1.41 | 1.34 ± 0.55 | ns |
| Lemongrass 1% | 1.24 ± 1.05 | 1.03 ± 0.69 | ns | 1.70 ± 1.26 | 0.96 ± 0.22 | 0.02 |
| **Picaridin 0.5%** | 2.44 ± 2.35 | 0.99 ± 0.57 | ns | 6.23 ± 9.77 | 1.15 ± 0.43 | ns |
| **Picaridin 1%** | 1.59 ± 1.20 | 1.39 ± 1.01 | ns | 1.55 ± 0.74 | 2.65 ± 2.96 | ns |
| **IR3535 0.5%** | 1.72 ± 1.58 | 1.16 ± 0.78 | ns | 1.61 ± 1.44 | 1.08 ± 0.54 | ns |
| **IR3535 1%** | 2.56 ± 1.38 | 1.64 ± 1.16 | 0.01 | 3.40 ± 3.50 | 0.96 ± 0.48 | 0.01 |

**Table 3.**

| Group | ISI 6<sup>th</sup> / ISI 1<sup>st</sup> |  | E.I. |
| --- | --- | --- | --- |
|  | MOP | OP |  |
| **Controls** | 0.479638009 | 0.576923077 | 1.2028 |
| **Eugenol 0.5%** | 1.105882353 | 0.604651163 | 0.5476 |
| **Eugenol 1%** | 0.586466165 | 0.4375 | 0.7459 |
| **Lemongrass 0.5%** | 0.768707483 | 0.623255814 | 0.8107 |
| **Lemongrass 1%** | 0.830645161 | 0.564705822 | 0.6798 |
| **Picaridin 0.5%** | 0.405737705 | 0.18459069 | 0.4549 |
| **Picaridin 1%** | 0.874213836 | 1.709677419 | 1.9556 |
| **IR3535 0.5%** | 0.674418605 | 0.670807453 | 0.9946 |
| **IR3535 1%** | 0.482352941 | 0.282352941 | 0.5853 |

## Discussion

### Effects of the repellents on the HOKN

Odor plumes are extremely important for insects’ foraging and exploration, and successful navigation relies on the multisensory integration of environmental and internal sensory inputs (Menti et al., 2023; Zjacic and Scholz, 2022). Attractive odours were reported for increasing the gain of gaze-stabilizing optomotor reflexes, maintaining the animal aligned within an invisible plume and facilitating the localization of attractive odorant sources while navigating in the environment (Frye, Tarsitano & Dickinson, 2003), and also for enhancing the final behavioural output through synergistical interaction with other sensory cues, like demonstrated for the visual, olfactory, and thermal receptors synergy in the landing response of *Anopheles coluzzii* mosquitoes (Carnaghi et al., 2021). However, in the evolutionary war between plants and insects, the former have been producing a variety of odorous-repellent substances or compounds that mask the odorous cues. And, although numerous studies have analysed the integration of attractive odorous stimuli with other sensory modalities in insects, mainly vision, relatively few studies are dedicated to investigating the interaction between olfactory and visual stimuli competing for one against the other. This becomes every day more important, in light of recent EU regulations pushing the use of natural aversive substances in pest control in agriculture to replace currently banned pesticides (Regulation (EU) No 528/2012). In this context, we observed that when a competing aversive odour is presented to flies undergoing optic flow stimulation (Table. 4):

1. the number of HOKN reduces, and their temporal distribution becomes more dispersed, reflecting an increment of the time interval separating consecutive events (ISI);
2. the velocity of the slow phase, associated with the movement tracking process of the flies, does not change overall.

**Table 4.**
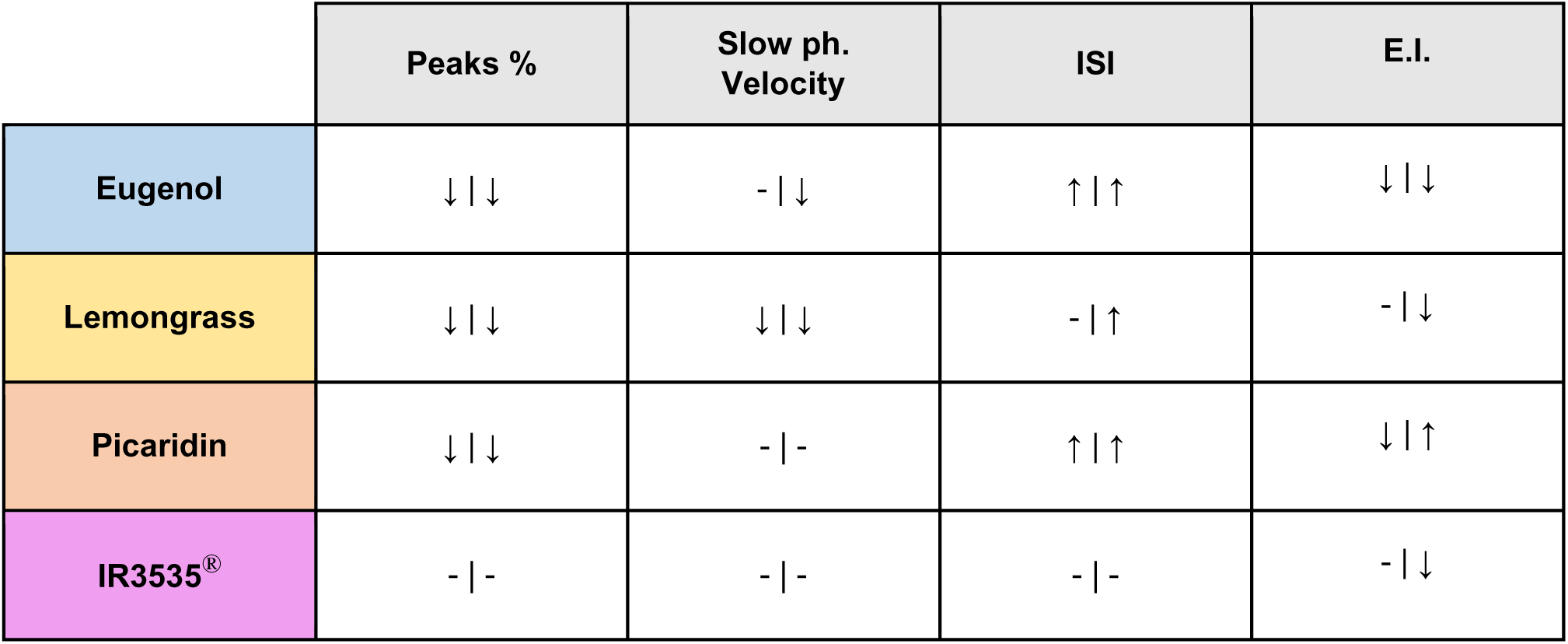

Additionally, we observed a slight decrease in the ISI as the trials proceeded, paired, in some of the groups, with an increment in the number of recorded HOKNs as a function of time.

Along with this general trend, a peculiar pattern emerges from the comparison between the repellent and masker sub-groups. As mentioned above, the substances chosen are classified as natural repellents (eugenol and lemongrass) or masking agents (IR3535^®^ and picaridin) for mosquitoes. In these insects, the two natural repellents strongly activate olfactory neurons, while the masking substances do not activate olfactory neurons substantially (Afify et al., 2019). Regarding *D. melanogaster*, lemongrass has been observed to be repellent in addition to being lethal as well after 96 hours of exposure (Aljedani, 2021; Yoon and Tak, 2022), and eugenol was shown to alter behaviour in adult animals and to also affect cardiac rhythm in larvae (Silva et al., 2022; Weineck et al., 2019). Considering, on the other hand, the two masking substances, the repellent action of these was also demonstrated in *Drosophila* with contrasting results: both IR3535^®^ and picaridin have shown repellent action in a choice inhibition test (Syed et al, 2011) but, in another experiment, the ‘masking’ effect of the former seemed more evident than a repellent action (Yoon and Tak, 2022).

Our experiments with *Drosophila* revealed that picaridin exhibits responses way closer to the natural repellents than to an expected masker as IR3535^®^ which in turn appears to have no effects on all the analysed parameters of the optokinetic response, as expected (Table 4). Taken together, our results show that repellents have an opposite action on the optomotor response compared to what was observed using attractors (Cheng et al., 2019), however, the mechanism of action seems to differ. Attractors can increase the gain of the optomotor response, improving it and resulting in an enhanced ability to localize resources (e.g. food). In the presence of a repellent, on the other hand, a decrease in the gain of the optomotor response is not observed (except in the higher concentration of lemongrass and with limited effects), but rather we do note a reduction in it in terms of jerks. In humans and mammals, a decrease in the gain of the optomotor response is associated with spatial disorientation: a reduction in nystagmus jerks is observed, for instance, in conditions of inattention, when individuals do not pay attention at all or only partially to the visual stimulus (Magnusson et al., 1985). It can then be assumed that the decrease in the number of jerks observed in the presence of repellents in flies may reduce the attentive ability and the resources allocated for localizing the source of the odour while maintaining the ability of the optomotor system to react promptly to perturbations. Moreover, a decreasing gain due to exposure to a repellent could underlie a possible, maybe dose-dependent, heavier effect if not toxicity of the compound (like the anaesthetic effect described in Weineck et al., 2019) which should then be avoided because of the fallout it could have on other sensitive insect species, like pollinators. Instead, synthetic maskers gave different results. IR3535^®^ did not affect both HOKN’s number and HOKN’s slow phase velocity. On the other hand, picaridin significantly reduced the number of HOKNs (MOP vs RP within the group), without altering the slow phase velocity. Therefore, in our study, IR3535^®^ appears to be neutral to the optomotor stimulus. Indeed, Yoon & Taak (2022) found a similar “masking” effect of IR3535^®^ in their experiments; however, conclusions are still unclear, as it has been observed that this compound elicits pb1 ORN-A neurons response in *Drosophila* (Syed et al., 2011), while calcium imaging does not reveal activation of antennal neurons in *Anopheles* after stimulation with the same compound, leaving doubts about how the substance exerts its physiological action. A different and more complex scenario is the outcome we observed after stimulation with picaridin. We found that picaridin significantly reduced the number of HOKNs. Similarly to IR3535^®^, picaridin was shown to activate pb1 ORN-A neurons in *Drosophila* (Syed et al., 2011), while also having a similar profile in *Anopheles* (Afify et al., 2019). Nevertheless, calcium imaging of antennal neurons showed some response in these cells after stimulation with picaridin (Fig. S3 in Afify et al., 2019), therefore suggesting that for both species picaridin may not be, after all, a neutral compound. Our data, obtained using a straightforward behavioural assay, agrees with these reported observations.

### Possible neurobiological mechanisms of the observed results

The visual information collected by the *ocelli* photoreceptors is processed in the optic lobe, which consists of the *lamina*, *medulla*, *lobula*, and *lobula* plate. Three distinct longitudinal channels of visual information processing (ON-OFF and colours) associated with horizontal analysis have been identified, which are able to distinguish the differences in movement and contrast between ommatidia (Borst et al., 2010). The visual information, processed at the level of the optic lobe, is then sent to three main nerve structures: the lateral cerebral ganglion, the central complex, and the mushroom bodies. The lateral cerebral ganglion (precisely in its PLP, PS, and PVLP components) is involved in the sensorimotor processing of decision-making processes, as it is the site of a multisensory convergence and the origin of numerous motor descending pathways (Namiki et al., 2018). Afferents to these regions from the tangential cells of the optic lobe (Borst et al., 2002) are involved in the processing of optomotor signals. Descending projection neurons, in addition to reaching the motor neuropile that innervates the indirect flight muscles and the leg muscles, also reach the motor neuropile for the neck muscles (Gronenberg and Strausfeld, 1990; Longden et al., 2022; Namiki et al., 2018; Strausfeld and Bassemir, 1985). Thus, through this pathway involving specific Descending Neurons (DN9), motor signals originating from optic lobe tangential cells and processing visual optic flow will eventually reach motor neurons regulating head movements (Frye and Dickinson, 2004; Namiki et al., 2018). Attractive odour plumes can increase the gain of optomotor response in *Drosophila*: This modulation of the visuomotor response is mediated by a specific *lobula* plate tangential interneuron, called Hx, which activity is enhanced whenever optic flow inputs are paired with odour inputs (Wasserman et al., 2015). Octopaminergic Tdc2 neurons, activated by odour inputs, are responsible for the modulation of Hx and of T4-T5 columnar neurons which in turn are also activated by optic flow inputs above tangential cells (Cheng et al., 2019; Wasserman et al., 2015). It was also suggested that T4/T5 neurons expressing octopaminergic receptors could be modulated by octopaminergic neurons and that this modulation can reverse the aversive effect of a visual object on fly behaviour (Cheng et al., 2019).

Our results seem to show a modulatory effect on the HOKN by repellents (eugenol, lemongrass, and picaridin to some extent) and no effects by masking odour plumes (IR3535^®^). Therefore, it is possible to assume that an octopaminergic modulation onto T4/T5 neurons and/or specific multimodal tangential cells (Hx tangential neuron) could take place when aversive compounds are delivered to the fly. Moreover, since DNs receive extensive dendritic contacts in the *lobula* plate, the aforementioned modulation may be reflected on the final motor output, thus possibly being also accountable for the alterations we found in the HOKNs.

*Learning may be altered by repellents but in an unsaturated environment*.

One last intriguing outcome of our experimental paradigm is the slight progressive increase in the number of saccades, and consequently reduced ISI duration, as the trials proceeded. To evaluate this aspect, we calculated the Effect Index (E.I.). Interestingly, the worst index values (i.e. most intrusive effect on the variable) resulted from the provenly effective repellents, but not for the Control and the IR3535 0.5% groups as expected. Notably, though, a quite low score of the index is indeed related to a greater concentration of IR3535^®^, which we did not expect. Moreover, in the Lemongrass group, the score is lower the higher the concentration, while the trend appears inverted in Eugenol and Picaridin (which exhibits both the most extreme scores in the table). This may suggest a possible effect of concentration, even in what we thought was an almost neutral compound from our other results (IR3535^®^), with maybe a possible lower saturation point for Eugenol and Picaridin with a consequent diminishing effect. Altogether, these results suggest both the possibility that flies can learn and improve their responsiveness to a prolonged constant optokinetic stimulation, and that being exposed to the repellents we tested could interfere, up to compound-specific saturation concentrations, with this process, effectively altering the flies’ learning dynamics in repellents unsaturated conditions.

In conclusion, we have shown here the application of a behavioural assay based on multisensory stimuli competition that can catch subtle differences in the action of aversive or neutral odorant compounds. This represents an ideal preliminary workbench to instruct more in-detail analyses, at the cellular or circuit level, to dissect the underlying neuronal mechanisms and potentially the impact of the compounds on the brain function.

## Materials and methods

### Flies’ strains

Berlin-K adult female flies aged 3-6 days were tested within 6 hours from the wake. Females were separated from males and collected under CO2 anaesthesia, then given at least 24 hours to recover.

The rearing was conducted in vials containing 10 ml of standard cornmeal medium, with a 12h:12h dark-light cycle and at room-controlled temperature (26 °C, at 45 ± 10% humidity).

Every group of flies was evaluated for only one (1) repellent and concentration, meaning that one group experienced, e.g., eugenol, concentrated at either 0.5% or 1%. The number of the analysed flies and total peaks, after pruning unsuccessful experiments and/or tracking (more in sections ahead), is reported in Fig. 4 (‘Results’ section).

### Fly’s preparation

Flies were transferred from the rearing vial to an empty one which was then put in ice. After the cold anesthetization, single flies (one at a time) were transferred to the mounting block, which was kept cold (+4°C) via a Peltier platform laid on a fan heat sink, and carefully placed upright inside the dedicated groove of the mounting support. The tip of a 34-gauge dispensing needle (BSTEAN, Shenzhen Hemasi E-Commerce Co., Ltd., PRC) was dipped in UV hard resin (DecorRom, Shenzhenshi Baishifuyou Trading Co., Ltd., PRC) removing the excess quantity (barely one drop remaining on the tip). The needle was then placed on one support angled at 60° and, with the aid of a micromanipulator, lowered onto the fly, touching the centre of the thorax; the resin was then cured for approximately 60 seconds with a UV torchlight to glue the animal to the pin. Flies were let to recover from the procedure for about 5-10 minutes; periodical small air puffs were delivered to verify the gluing had been performed correctly, as well as the willingness of the animal to fly. Badly glued or unwilling flies were discarded. Once the fly was glued and actively flying, the pin was transferred and mounted inside the experimental setup.

### Experimental apparatus

The experiment is conducted in a home-built dark chamber (components bought from Thorlabs Inc.), with side access, containing all the hardware.

A vertical support, ending with a syringe attachment, where the pin together with the glued fly can be secured, is suspended above the camera (Q4500, Ximea GmbH, Germany) equipped with an infra-red (IR) bandpass filter; two IR LEDs (Thorlabs Inc., US) provide the necessary illumination.

The other components of the apparatus include:

- an adjustable hand-crafted screen, made of parchment paper, placeable in front of the animal, covering 180° around the fly;
- the projector (Lightcrafter 4500, Texas Instruments Inc.), placed in front of the screen;
- the custom odour-<u>delivery</u> system made up of plastic tubing of various diameters, assorted luers (Ark-Plas Products Inc., US), glass capillaries (GB150F-10, Science Products GmbH, Germany), an Arduino (UNO REV3, Arduino, US) controlled solenoid valve (SIRAI Elettromeccanica S.r.l., Italy), two glass vials containing the solutions, and an air pump (Air Professional 150, PRO.D.AC INTERNATIONAL S.r.l., Italy) for the vaporizing;
- the custom odour-<u>recycling</u> system, consisting of plastic tubing, an externally alimented suction unit (VN-C4 vacuum pump, You Cheng Industrial Co., Ltd., Taiwan), plus the glass flask used to achieve negative pressure and the subsequent suction.

Due to computer specs limitations, we had to use two different computers, one for running the protocol while the second took care of the recording. The two machines were made to communicate via a U3-LV Labjack (Labjack Corporation, US): the recording of the experiment, controlled by the first computer, would only start after receiving the signal sent from the second computer when starting the protocol through the dedicated script.

We wrote down custom scripts in MATLAB, to control the camera and the recording, and in Python, to control the presentation of the visual stimuli (designed through the open-source PsychoPy© toolbox, Open Science Tools Ltd., UK), the modulation of the solenoid valve, and also for synchronizing the start between the protocol script and the recording.

Due to the utilization of two different computers and the MATLAB-mediated camera control, the framerate of acquisition could not exceed 45 frames per second on average.

### Repellent compounds

The chosen repellent substances (eugenol, lemongrass oil, picaridin, and IR3535® *alias* ‘Nb[n-N-butyl-N-acetyl] aminopropionic acid ethyl ester’) were purchased at the highest purity available from Biosynth® Ltd, UK.

### Experimental paradigm

The paradigm was structured in 6 (six) repetitions of one trial (Fig. 2), where the flies faced optokinetic stimulation both in the absence and presence of the repellent odour plume. The trial was structured as follows:

*a.* one 15° vertical black bar on a white background, presented in the middle of the screen (duration: 10 seconds)
*b.* a mask of gratings made up of 15° alternate black and white vertical bars moving clockwise at a fixed speed of 60 deg.sec^-1^ (duration: 60 seconds)
*c.* same as (*a.*)
*d.* pause in darkness (duration: 20 seconds)

The air plume was continuous for the duration of the trials and coming from the right to the left of the animal, or in the direction opposite to the optokinetic stimulation (Fig. 1A). Every time the gratings phase (b.) reached its half (30 seconds), the solenoid valve was switched through the Arduino board, as the pre-defined switch state was properly set to ‘ON’ from within the Python script. The activation of the valve switched the air intake and output from the odourless vial (pure mineral oil, MOP phase) to the repellent one (compound in solution with mineral oil, OP phase), so that, in every trial, the flies experienced 30 seconds of visual stimulation within an odour neutral air flow followed by another 30 seconds of the same visual stimulus but in presence of the competing odorous compound.

### Data extraction

The tracking of the animal was conducted offline through the self-contained MATLAB program ‘Flyalyzer’ (Rauscher and Fox, 2021). Bad tracking results, which could be caused by illumination issues and/or excessive leg movements from the fly’s self-grooming interfering with the tagging of the antennae, were discarded.

From the obtained raw data of the head movements, we wrote a custom MATLAB script exploiting a native function (*find peaks*) to extract a template of movement congruent with the optomotor response. This template was then utilized to identify the peaks on the track corresponding to the optomotor events (Fig. 1C and Fig. 2). For further clarification: it is known that flies also perform a small fraction of co-directional saccades when presented with moving gratings (Cellini et al., 2021), therefore, the raw head tracks contained both positive and negative peaks. Since we were not interested in the co-directional saccades, we wrote the script to only tag the (positive, in our coordinates system) head-reset fast movements and the related slow phases.

Subsequent data elaboration, plotting, and statistical analysis were performed in RStudio through custom-made scripts.

### Statistical analysis

The whole analysis was conducted with α = 0.95.

Data were checked for normality (Shapiro-Wilkins test) and heteroscedasticity (Bartlett’s test when the distributions were not normal) across groups.

The statistical relevance of the observed differences in the number of events (HOKN) was assessed through paired Z-tests with continuity correction, checking if the proportion of saccades identified during the MOP differed significantly with respect to the OP between the “Controls” group and the other compounds plus within same compounds at different concentrations.

ISI data were not normally distributed nor homogenous in variances. However, given the large amount of data, we analysed the ISI distribution opting for a 2-Way ANOVA (accounting for the presence/absence of the compound during the OKR task and the groups of repellent tested) corrected for the non-heteroscedasticity, followed by a post-hoc Games-Howell test corrected for non-normality of the data.

Confrontations between the 1^st^ and 6^th^ trials ISI (both MOP and OP), as well as the slow-phase velocity profiles, were performed for each group through the Mann-Whitney U test for paired data, as data were not normally distributed.

## Abbreviations

C: control group
E05: group exposed to eugenol 0.5% solution
E1: group exposed to eugenol 1% solution
HOKN: head optokinetic nystagmus
I05: group exposed to IR3535® 0.5% solution
I1: group exposed to IR3535® 1% solution
L05: group exposed to lemongrass 0.5% solution
L1: group exposed to lemongrass 1% solution
IR: infra-red
ISI: inter-saccade interval
MOP: mineral oil phase
OF: optic flow
OKN: optokinetic nystagmus
OKR: optokinetic response
OP: odorant phase
P05: group exposed to picaridin 0.5% solution
P1: group exposed to picaridin 1% solution.

## Data Availability

An R-markdown file retracing the content of the paper, and the relevant produced dataset, are available in the “HOKN-Flies” repository, at <u>Github.com</u>; in addition the pre-processed data, along with a video sample are hosted in the “Optokinetic response in D. Melanogaster reveals the nature of common repellent odorants” repository at Zenodo.com (DOI: 10.5281/zenodo.11183967).

The Arduino and Python scripts, related to the experimental paradigm control, and the scripts written for the data analysis (mostly included in the markdown document), can be shared by the authors upon motivated request.

## Acknowledgements

Authors thank Claudia Lodovichi, Marco Brondi, and Antonio Di Soccio, for their constructive criticism and suggestions regarding the construction of the home-made odour-delivery system; Maria Elena Miletto-Petrazzini for her support in the experimental paradigm conceptualization and designing.

## Authors contribution

GMM, PV, AD, and AM designed the experiment. GMM, MB, MAZ, and AM, set up the experimental apparatus and wrote the related scripts. GMM designed the final experimental paradigm and acquired the data. GMM, MAZ, and AM carried out the pre-processing and data analysis. GMM drafted the manuscript and prepared the figures and tables, AM and MDM drafted the Discussion section. All authors contributed to the interpretation of data and the revision of the manuscript.

## References

Afify A. et al. (2019) Commonly Used Insect Repellents Hide Human Odors from Anopheles Mosquitoes. Current Biology 29 (21): 3669–3680. 10.1016/j.cub.2019.09.007

Aljedani D. M. (2021) Effects of Some Insecticides (Deltamethrin and Malathion) and Lemongrass Oil on Fruit Fly (*Drosophila melanogaster*). Pak. J. Biol. Sci. 24, 477–491. 10.3923/pjbs.2021.477.491

Borst A., Haag J. (2002) Neural networks in the cockpit of the fly. J. Comp. Physiol. A 188, 419–437. 10.1007/s00359-002-0316-8

Borst A., Haag J. & Reiff D. F. (2010) Fly Motion Vision. Annual Review of Neuroscience vol. 33 49–70. 10.1146/annurev-neuro-060909-153155

Cellini B. et al. (2021) Mechanisms of punctuated vision in fly flight. Curr. Biol. 31: 4009–4024 10.1016/j.cub.2021.06.080

Cheng K. Y., Colbath R. A., and Frye M. A. (2019). Olfactory and Neuromodulatory Signals Reverse Visual Object Avoidance to Approach in Drosophila. Curr. Biol. 29: 2058–2065.e2. 10.1016/j.cub.2019.05.010

Chow D. M., Frye M. A. (2009) The neuro-ecology of resource localization in Drosophila: behavioral components of perception and search. Fly (Austin) 3: 50–61. 10.4161/fly.3.1.7775

Chow D. M., Frye M. A. (2008) Context-dependent olfactory enhancement of optomotor flight control in Drosophila. J. Exp. Biol. 211 (15): 2478–2485. 10.1242/jeb.018879

Carnaghi M., Belmain S. R., Hopkins R. J., Hawkes F. M. (2021) Multimodal synergisms in host stimuli drive landing response in malaria mosquitoes. Sci. Rep. 11, 7379. 10.1038/s41598-021-86772-4

Currier T. A., Nagel K. I. (2020) Multisensory control of navigation in the fruit fly. Current Opinion in Neurobiology 64: 10–16. 10.1016/j.conb.2019.11.017

Duistermars B. J. and Frye M. A. (2010) Multisensory integration for odor tracking by flying Drosophila. Communicative & Integrative Biology, 3 (1): 60–63. 10.4161/cib.3.1.10076

Duistermars B. J., Chow D. M., Frye M. A. (2009) Flies require bilateral sensory input to track odor gradients in flight. Current Biology 19: 1301–1307. 10.1016/j.cub.2009.06.022

Farries M. A. (2013) How “basal” are the basal ganglia? Brain, Behavior and Evolution 82: 211–214. 10.1159/000356101

Frye M. A., Tarsitano M. & Dickinson M. H. (2003) Odor localization requires visual feedback during free flight in Drosophila melanogaster. J. Exp. Biol. 206, 843–855. 10.1242/jeb.00175

Frye M. A., Dickinson M. H. (2004) Motor output reflects the linear superposition of visual and olfactory inputs in Drosophila. J. Exp. Biol. 207 (1): 123–131. 10.1242/jeb.00725

Götz K.G. (1964) Optomotorische Untersuchung des visuellen systems einiger Augenmutanten der Fruchtfliege Drosophila. Kybernetik 2: 77–92. 10.1007/BF00288561

Giunti G., et al. (2023) Invasive mosquito vectors in Europe: From bioecology to surveillance and management. Acta Tropica 239. 10.1016/j.actatropica.2023.106832

Gronenberg W. and Strausfeld, N. J. (1990) Descending neurons supplying the neck and flight motor of diptera: Physiological and anatomical characteristics. J. Comp. Neurol., 302: 973–991. 10.1002/cne.903020420

Hecht S., Wald G. (1934) The visual acuity and intensity discrimination of Drosophila. J Gen Physiol 17 (4): 517–547. 10.1085/jgp.17.4.517

Land M. (2019) Eye movements in man and other animals. Vision Research 162: 1–7. 10.1016/j.visres.2019.06.004

Longden K. D., Schützenberger A., Hardcastle B. J. & Krapp H. G. (2022) Impact of walking speed and motion adaptation on optokinetic nystagmus-like head movements in the blowfly Calliphora. Ski. Rep. 12, 11540. 10.1038/s41598-022-15740-3

Magnusson, M., Pyykkö, I. & Jäntti, V. (1985) Effect of alertness and visual attention on optokinetic nystagmus in humans. American Journal of Otolaryngology 6, 419–425. 10.1016/S0196-0709(85)80020-X

Moroz L. L. (2009) On the independent origins of complex brains and neurons. Brain, Behavior and Evolution 74: 177–190. 10.1159/000258665

Namiki, S., Dickinson, M. H., Wong, A. M., Korff, W. & Card, G. M. (2018) The functional organization of descending sensory-motor pathways in Drosophila. eLife 7, e34272. 10.7554/eLife.34272

Northcutt R. G. (2012) Evolution of centralized nervous systems: two schools of evolutionary thought. Proceedings of the National Academy of Sciences of the United States of America 109 (Suppl. 1): 10626–10633. 10.1073/pnas.1201889109

Rauscher M. J. and Fox J. L. (2021) Haltere and visual inputs sum linearly to predict wing (but not gaze) motor output in tethered flying Drosophila. Proceedings of the Royal Society B. 2882020237420202374.

10.1098/rspb.2020.2374

Semenza J. C. and Suk J.E. (2018), Vector-borne diseases and climate change: a European perspective, FEMS Microbiology Letters, 365 (2): 244. 10.1093/femsle/fnx244

Silva, J. C. et al. (2022) Evaluation of antibacterial and toxicological activities of essential oil of Ocimum gratissimum L. and its major constituent eugenol. Food Bioscience 50, 102128. 10.1016/j.fbio.2022.102128

Strausfeld N. J. & Bassemir U. K. (1985) Lobula plate and ocellar interneurons converge onto a cluster of descending neurons leading to neck and leg motor neuropil in Calliphora erythrocephala. Cell Tissue Res. 240, 617–640. 10.1007/BF00216351

Strausfeld N. J. and Hirth F. (2013a) Homology versus convergence in resolving transphyletic correspondences of brain organization. Brain Behav. Evol. 82: 215–219. 10.1159/000356102

Strausfeld N. J. and Hirth F. (2013b) Deep Homology of Arthropod Central Complex and Vertebrate Basal Ganglia. Science 340: 157–161. 10.1126/science.1231828

Syed Z., et al. (2011) Generic Insect Repellent Detector from the Fruit Fly Drosophila melanogaster. PLoS ONE 6 (3): e17705. 10.1371/journal.pone.0017705

Tait G. et al (2021), Drosophila suzukii (Diptera: Drosophilidae): A Decade of Research Towards a Sustainable Integrated Pest Management Program, Journal of Economic Entomology 114, Issue **5**: 1950–1974. 10.1093/jee/toab158

Wasserman, S. M. et al. (2015) Olfactory Neuromodulation of Motion Vision Circuitry in Drosophila. Current Biology 25: 467–472. 10.1016/j.cub.2014.12.012

Weineck K, Stanback A, Cooper RL. (2019) The Effects of Eugenol as an Anesthetic for an Insect: Drosophila, Adults, **Larval Heart Rate, and Synaptic Transmission**. Proceedings of the Association for Biology Laboratory Education, 40: 54. https://web.as.uky.edu/biology/faculty/cooper/labWWW-PDFs/ABLE-Fly-Eugenol%20paper-2019.pdf

Yoon, J. & Tak, J. H. (2022) Toxicity and context-dependent repellency of temporarily granted repellents under new biocidal products regulations in South Korea against Drosophila melanogaster (Diptera: Drosophilidae). J. Asia-Pac. Entomol. 25, 2, 101911. 10.1016/j.aspen.2022.101911

Zjacic, N., and Scholz, M. (2022). **The role of food odor in invertebrate foraging**. Genes, Brain Behav. 21, e12793. 10.1111/gbb.12793

## Online material

1. FAO/IAEA, Guilhot R., Taret G., Gembinsky K. and Cáceres C. (eds.) (2022). **Guidelines for Mass Rearing and Irradiation of Drosophila suzukii for Sterile Insect Technique Application**., Food and Agriculture Organization of the United Nations/International Atomic Energy Agency. Vienna, Austria. https://www.iaea.org/sites/default/files/massrearing-and-irradiation-swd.pdf

2. FAO/IAEA (2021) Guidelines for Biosafety and Biosecurity in Mosquito Rearing Facilities. Food and Agriculture Organization of the United Nations/International Atomic Energy Agency. Vienna, Austria https://www.iaea.org/sites/default/files/guidelines_for_mosquito_facilities.pdf

3. **Regulation (EU) No 528/**2012 of the European Parliament and of the Council of 22 May 2012 concerning the making available on the market and use of biocidal products. http://data.europa.eu/eli/reg/2012/528/2022-04-15

