## Supplementary Figures and Table for "Optokinetic response in *D. Melanogaster* reveals the nature of common repellent odorants"

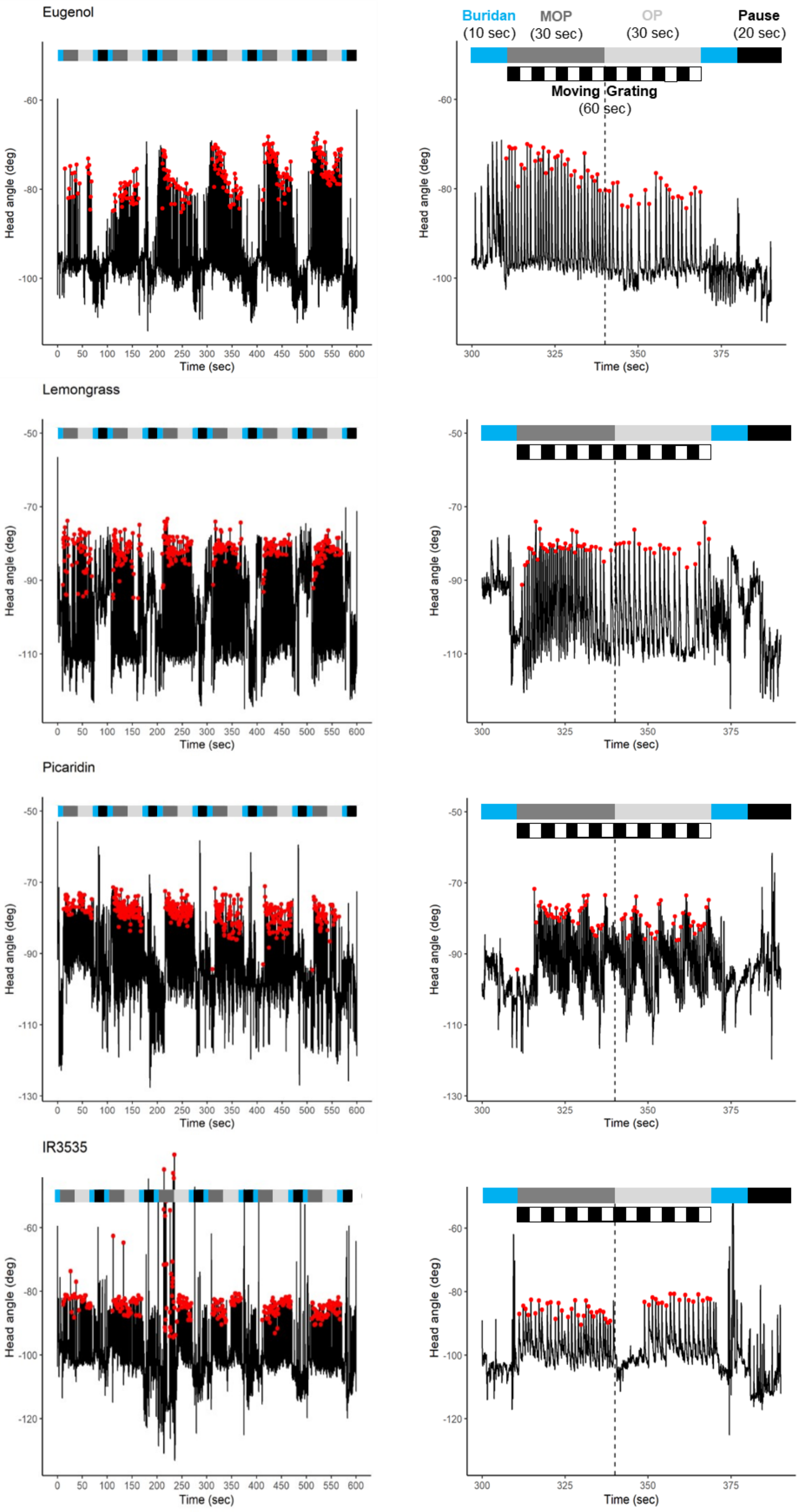


**Figure 1. Raw Tracks.** Raw tracks with tagged HOKNs and 4th trial zoom, as shown in the main text.

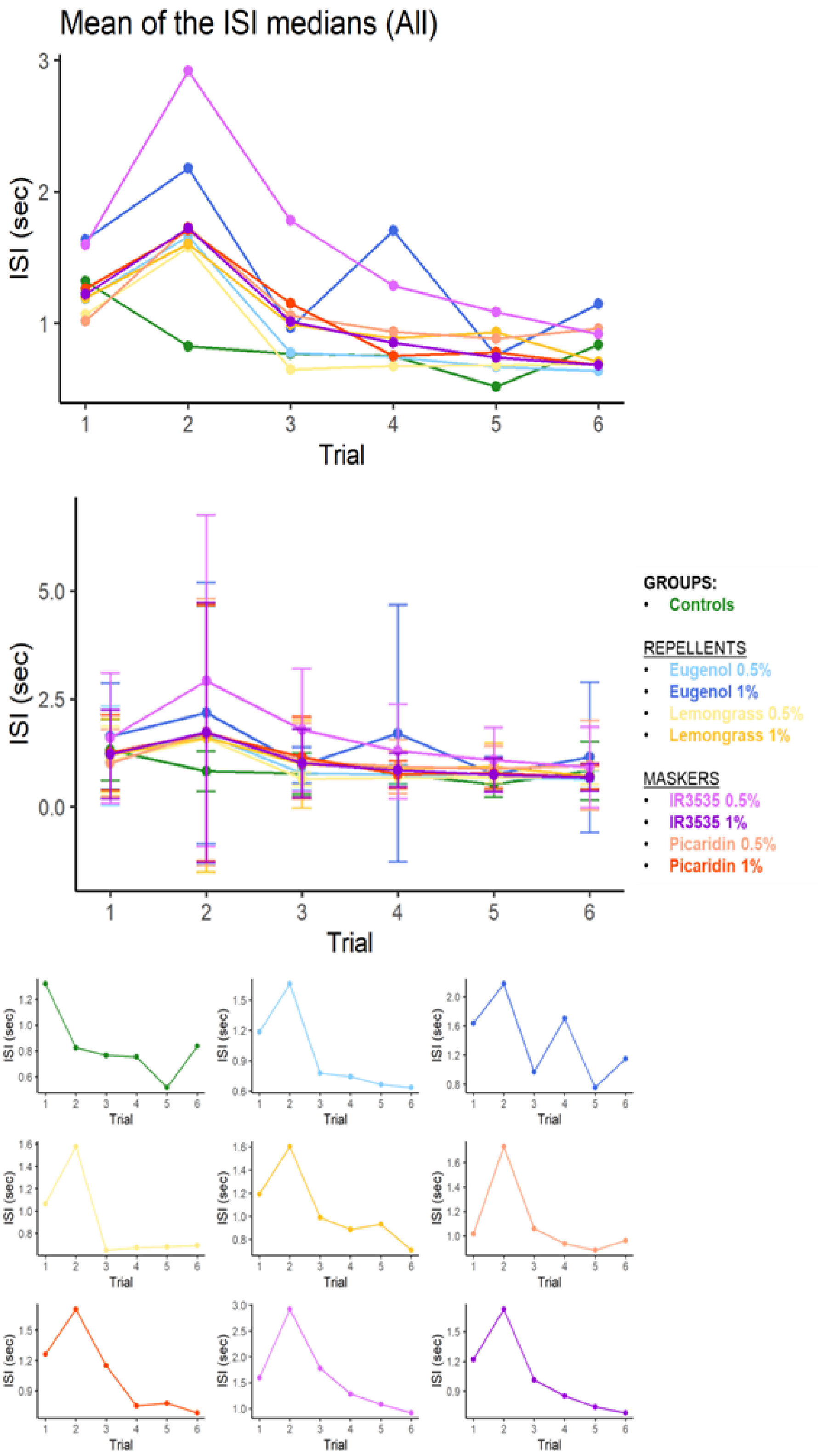


**Figure 2. ISI across all trials.** Mean among subjects of the median ISI value across trials, plotted together without and with the standard deviation. In the last plot groups are plotted alone.

| | Comparison of proportion of peaks over time | | | | | | | |
|---|---|---|---|---|---|---|---|---|
| | Controls | Eugenol | Eugenol05 | IR3535 | IR353505 | Lemongrass | Lemongrass05 | Picaridin |
| Eugenol | 13.23769 | | | | | | | |
| | 0.0000* | | | | | | | |
| Eugenol05 | 11.43379 | -1.803900 | | | | | | |
| | 0.0000* | 1.0000 | | | | | | |
| IR3535 | 12.01751 | -1.201626 | 0.599745 | | | | | |
| | 0.0000* | 1.0000 | 1.0000 | | | | | |
| IR353505 | -1.759253 | -15.00617 | -13.20101 | -13.78265 | | | | |
| | 1.0000 | 0.0000* | 0.0000* | 0.0000* | | | | |
| Lemongrass | 2.026688 | -11.22023 | -9.415075 | -10.00202 | 3.788585 | | | |
| | 0.7685 | 0.0000* | 0.0000* | 0.0000* | 0.0027* | | | |
| Lemongrass05 | 6.607724 | -6.629969 | -4.826069 | -5.419051 | 8.371584 | 4.585641 | | |
| | 0.0000* | 0.0000* | 0.0000* | 0.0000* | 0.0000* | 0.0001* | | |
| Picaridin | 9.062556 | -4.193561 | -2.387150 | -2.984373 | 10.83059 | 7.039378 | 2.445636 | |
| | 0.0000* | 0.0005* | 0.3056 | 0.511 | 0.0000* | 0.0000* | 0.2603 | |
| Picaridin05 | 5.500115 | -7.709699 | -5.909598 | -6.499836 | 7.259477 | 3.481519 | -1.093692 | -3.535679 |
| | 0.0000* | 0.0000* | 0.0000* | 0.0000* | 0.0000* | 0.0090* | 1.0000 | 0.0073* |

**Supplementary Table 1. Dunnet test.** Check on the differences in the distribution of HOKNs over time.
